# Crystal structure of chloroplastic thioredoxin z defines a novel type-specific target recognition

**DOI:** 10.1101/2020.11.24.396572

**Authors:** Théo Le Moigne, Libero Gurrieri, Pierre Crozet, Christophe H. Marchand, Mirko Zaffagnini, Francesca Sparla, Stéphane D. Lemaire, Julien Henri

**Affiliations:** Sorbonne Université, CNRS, UMR 7238, Institut de Biologie Paris-Seine, Laboratoire de Biologie Computationnelle et Quantitative, 4 place Jussieu F-75005 Paris, France; Faculty of Sciences, Doctoral School of Plant Sciences, Université Paris-Saclay, F-91190 Saint-Aubin, France; Sorbonne Université, CNRS, UMR 8226, Institut de Biologie Physico-Chimique, Laboratoire de Biologie Moléculaire et Cellulaire des Eucaryotes, 13 rue Pierre et Marie Curie F-75005 Paris, France; Department of Pharmacy and Biotechnology, University of Bologna, via Irnerio 42, I-40126 Bologna, Italy; Sorbonne Université, Polytech Sorbonne, F-75005 Paris, France; CNRS, Institut de Biologie Physico-Chimique, Plateforme de Protéomique, FR 550, 13 rue Pierre et Marie Curie F-75005 Paris, France

## Abstract

Thioredoxins (TRXs) are ubiquitous disulfide oxidoreductases structured according to a highly conserved fold. TRXs are involved in a myriad of different processes through a common chemical mechanism. Plant thioredoxins evolved into seven types with diverse subcellular localization and distinct protein targets selectivity. Five TRX types coexist in the chloroplast, with yet scarcely described specificities. We solved the first crystal structure of a chloroplastic z-type TRX, revealing a conserved TRX fold with an original electrostatic surface potential surrounding the redox site. This recognition surface is distinct from all other known TRX types from plant and non-plant sources and is exclusively conserved in plant z-type TRXs. We show that this electronegative surface endows TRXz with a capacity to activate the photosynthetic Calvin-Benson cycle enzyme phosphoribulokinase. TRXz distinct electronegative surface thereby extends the repertoire of TRX-target recognitions.

## Introduction

Thioredoxins (TRXs) are small ubiquitous disulfide oxidoreductases [1–5]. TRXs fold into a highly conserved and thermostable domain composed of a mixed β-sheet closely surrounded by α-helices and exposing a WC(G/P)PC pentapeptidic motif [3, 6]. Phylogenetic analyses described the history of the TRX fold through four billion years of evolution into the contemporary proteins [7, 8]. TRXs modify the ternary or quaternary structures of proteins by reducing target disulfide bonds into separate thiols [9, 10]. TRXs have also been proposed to control other redox modifications of cysteines by catalyzing denitrosylation, deglutathionylation, or depersulfidation reactions, contributing therefore to redox signaling cascades and metabolic remodeling [11–15], as recently reviewed in [16, 17]. TRXs midpoint redox potential is comprised between −310 and −230 mV at pH 7 [18–22] with a nucleophilic active site cysteine reacting in the thiolate state (–S^−^), favored at physiological pH by a local environment determining a cysteine p*K*_a_ in the 6.5-7.5 range [23–25].

The TRX system is recognized as having multiple roles in a myriad of cellular processes and numerous human diseases [1, 26–28]. Non-photosynthetic organisms contain a limited number of TRXs reduced by NADPH:thioredoxin reductase (NTR). By contrast, TRXs form a larger multigene family in photosynthetic organisms (21 isoforms in the model plant *Arabidopsis thaliana* and ten in the unicellular green alga *Chlamydomonas reinhardtii*). Phylogenetic analyses grouped TRXs in cytosolic/mitochondrial h-type, mitochondrial o-type and five chloroplastic types (f-, m-, x-, y- and z-types) [29, 30]. Cytosolic and mitochondrial TRXs are reduced by NTR but chloroplastic TRXs are specifically reduced in the light by ferredoxin:thioredoxin reductase (FTR) which derives electrons from ferredoxin and the photosynthetic electron transfer chain [16, 30–34]. This unique reduction mechanism allows to couple the redox state of TRX to light intensity and to use the chloroplast TRX system as a light-dependent signaling pathway for regulation of cellular metabolism and processes [35]. In photoautotrophic organisms, TRXs were originally identified as light-dependent regulators of the Calvin-Benson cycle (CBC), responsible for fixation of atmospheric CO_2_ into triose phosphates using energy (ATP) and reducing power (NADPH) produced in the light by the photosynthetic electron transfer chain. Early studies identified four key enzymes of the CBC as TRX targets: phosphoribulokinase (PRK) [36], glyceraldehyde-3-phosphate dehydrogenase (GAPDH) [37], fructose-1,6-bisphosphatase (FBPase) [38] and sedoheptulose-1,7-bisphosphatase (SBPase) [39]. These enzymes have a low activity in the dark and are activated upon illumination by the ferredoxin/TRX system because reduction of specific regulatory disulfides by TRX triggers conformational changes that shift the enzyme to a high activity conformation. Besides the CBC, chloroplast TRXs were later recognized as regulators of multiple targets involved in numerous pathways and processes [16, 40]. TRXs were especially recognized to provide electrons for the regeneration of major antioxidant enzymes such as peroxiredoxins [21, 41–46].

In the microalga *Chlamydomonas reinhardtii* ten TRX isoforms have been identified including six nuclear-encoded chloroplastic TRXs: CrTRXf1, CrTRXf2, CrTRXm, CrTRXx, CrTRXy, and CrTRXz [16, 30, 40]. Proteomic analyses based on affinity purification chromatography and *in vitro* reconstitution of the cytosolic TRX system allowed the identification of 1053 TRX targets and 1052 putative regulatory sites in Chlamydomonas [35, 47]. Among these, all CBC enzymes were identified indicating that they are potential TRX interactors, and direct TRX-dependent activation of CBC enzymes was only demonstrated for PRK [48], FBPase [49], SBPase [50] and phosphoglycerate kinase (PGK) [51]. By contrast with land plants, photosynthetic GAPDH from *Chlamydomonas reinhardtii* (CrGAPDH) is not directly regulated by TRX but only indirectly through formation of the (A_4_-GAPDH)_2_-CP12_4_-PRK_2_ complex [52–54]. Among the five TRX types present in chloroplasts, only TRXf has been systematically compared to other types and demonstrated to target FBPase [21, 32], PRK [48, 54], GAPDH in land plants [54] and PGK in Chlamydomonas [51]. Structural analysis revealed that electrostatic complementarity was the principal driver of TRXf-targets recognition [55–59]. Besides, TRXs were recently attributed a reciprocal redox function for the oxidation of targets upon light to dark transitions through the action of TrxL2 and 2-Cys peroxiredoxins [41, 60–62]. Finally, TRXs are involved in a multiplicity of other plastid functions either through their redox capacities [34], in the complex redox cellular network [63, 64] or through participation in large supramolecular assemblies [65, 66].

In *Populus trichocarpa* and in *Arabidopsis thaliana*, TRXz was reported to be reduced by FTR [67, 68]. TRXz can also act as a potential electron acceptor for a special type of chloroplastic TRX named NADPH:thioredoxin reductase C (NTRC) [64], TRXf, m, x and y [69], and interacts with fructokinase-like proteins (FLN1 and FLN2) in *Arabidopsis thaliana* and *Nicotiana benthamiana* [70]. TRXz-FLN interaction regulates plastid-encoded polymerase (PEP) [70, 71]. Notably, TRXz reduces plastid redox insensitive protein 2 (PRIN2) cysteine 68 (C68) hetero-molecular disulfides bridge within a homodimer in *Arabidopsis thaliana*, releasing monomeric PRIN2 that activates PEP transcription [72]. TRXz redox activity may however be dispensable and TRXz-FLN1 are proposed to be essential components of the PEP complex [73]. TRXz also interacts with *Arabidopsis thaliana* thioredoxin-like MRL7 [74] and Temperature-Sensitive Virescent (TSV) protein mediates TRXz interaction with PEP in *Oryza sativa* [75]. In rice, TRXz was also found to interact with chloroplastic RNA editing enzymes [76]. Yet, the full set of TRXz targets or interacting partners is still unknown and the corresponding molecular basis for TRXz specificity towards its targets has to be deciphered.

Altogether, it is striking that the very ancient and simple 12-15 kDa TRX fold has evolved into this plethora of functions, suggesting a fine selection of redox properties and specific surface recognitions. In order to gain insights into the molecular functions of chloroplastic z-type TRXz and the physico-chemical basis for its specificity, we solved the high-resolution crystal structure of TRXz from *Chlamydomonas reinhardtii*. This structure is the first representative of a z-type TRX and unravels a conserved native folding compared to other structurally solved TRXs. Model analysis mapped its redox site and identified the surfaces it exposes for selective protein recognition. TRXz displays electro-complementary surfaces with CrPRK. We show that PRK is indeed activated by TRXz reduction *in vitro*.

## Results

### CrTRXz folds as a canonical thioredoxin

Recombinant TRXz from *Chlamydomonas reinhardtii* (CrTRXz) was heterologously expressed in *E. coli* as a 149 amino acids polypeptide (predicted mature protein including residues 56-183 plus the N-terminal affinity tag) and purified to homogeneity by metal-affinity chromatography. Purified CrTRXz was crystallized, the crystals were cryoprotected and submitted to X-ray diffraction for the collection of a complete dataset indexed in space group P3_2_21 and solved by molecular replacement with CrTRXf2 (PDB ID code: 6I1C) as a search model. Model correction and completion up to 116 amino acid residues and one water molecule was refined to R=0.2296 and R_free_=0.2335 at a resolution of 2.4 Å (Table 1). CrTRXz folds according to the canonical TRX topology (SCOPe entry c.47), *i*.*e*. a central mixed β-sheet of four strands sandwiched between two pairs of α-helices (Figure 1A and B). The strands order is 2-1-3-4 with strand 3 antiparallel to strands 1, 2 and 4. The succession of secondary structures is as follows: residues 67-76 form helix 1, residues 80-86 strand 1, residues 91-107 helix 2, residues 111-117 strand 2, residues 122-128 helix 3, residues 135-139 strand 3, residues 148-151 strand 4 and residues 156-167 helix 4 (Figure S1). Length of helix 1 is conserved in the eukaryotic branch of TRX evolution since the last eukaryotic common ancestor [7]. Residues 118-120 were modelled as one turn of a helix 3_10_ with D119 hydrogen bonding with W89 and the unique water molecule of the model. Residues ^54^HMVI^57^ and ^61^KVEKIS^66^ of recombinant CrTRXz form an unfolded extension at the amino-terminus of the model, deforming it from an ideal sphere. Residues 58-60 and 170-183 were not built because of a lack of interpretable electron density. Pairwise alignments of equivalent C_α_ of CrTRXf2 and CrTRXm onto CrTRXz yielded a root mean square deviation (RMSD) of 0.838 Å and 1.356 Å respectively, confirming the close structural similarity between the experimentally solved Chlamydomonas TRX structures (Figure S2). Further alignments with experimentally determined structures from the protein data bank (PDB) revealed also high similarities with *Plasmodium falciparum* TRX2 (PDB identifier 3UL3, RMSD=1.16 Å), resurrected ancestral TRX from inferred last bacterial common ancestor (4BA7, RMSD=1.18 Å), *Bacteroides fragilis* TRXP (3HXS, RMSD=1.19 Å), and with 20 other TRXs with RMSD ≤ 1.30 Å. CrTRXz topology is classical among TRXs.

**Table 1.**
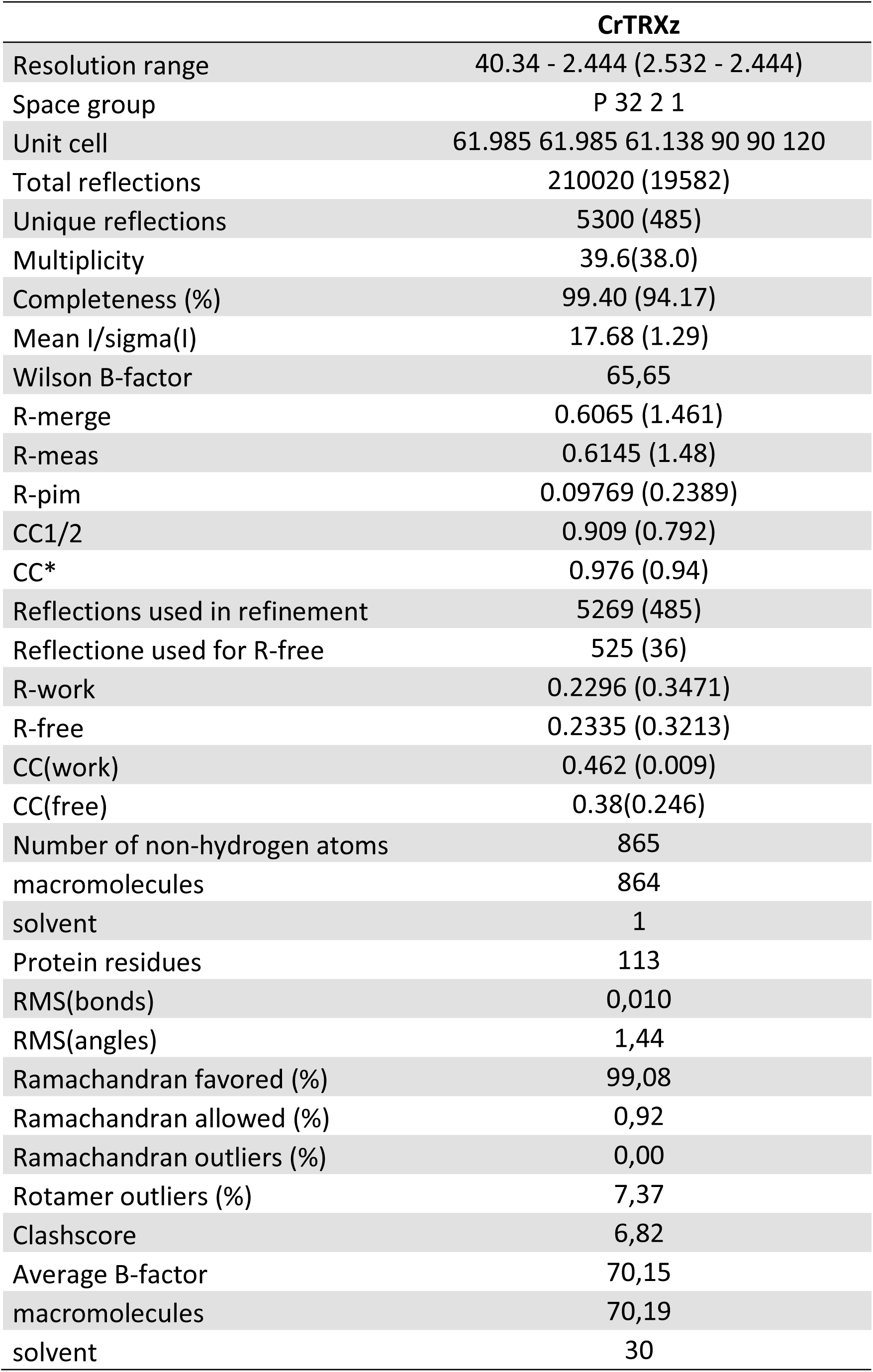
Crystallographic diffraction data and model statistics. Statistics for the highest-resolution shell are reported in parentheses.

**Figure 1.**
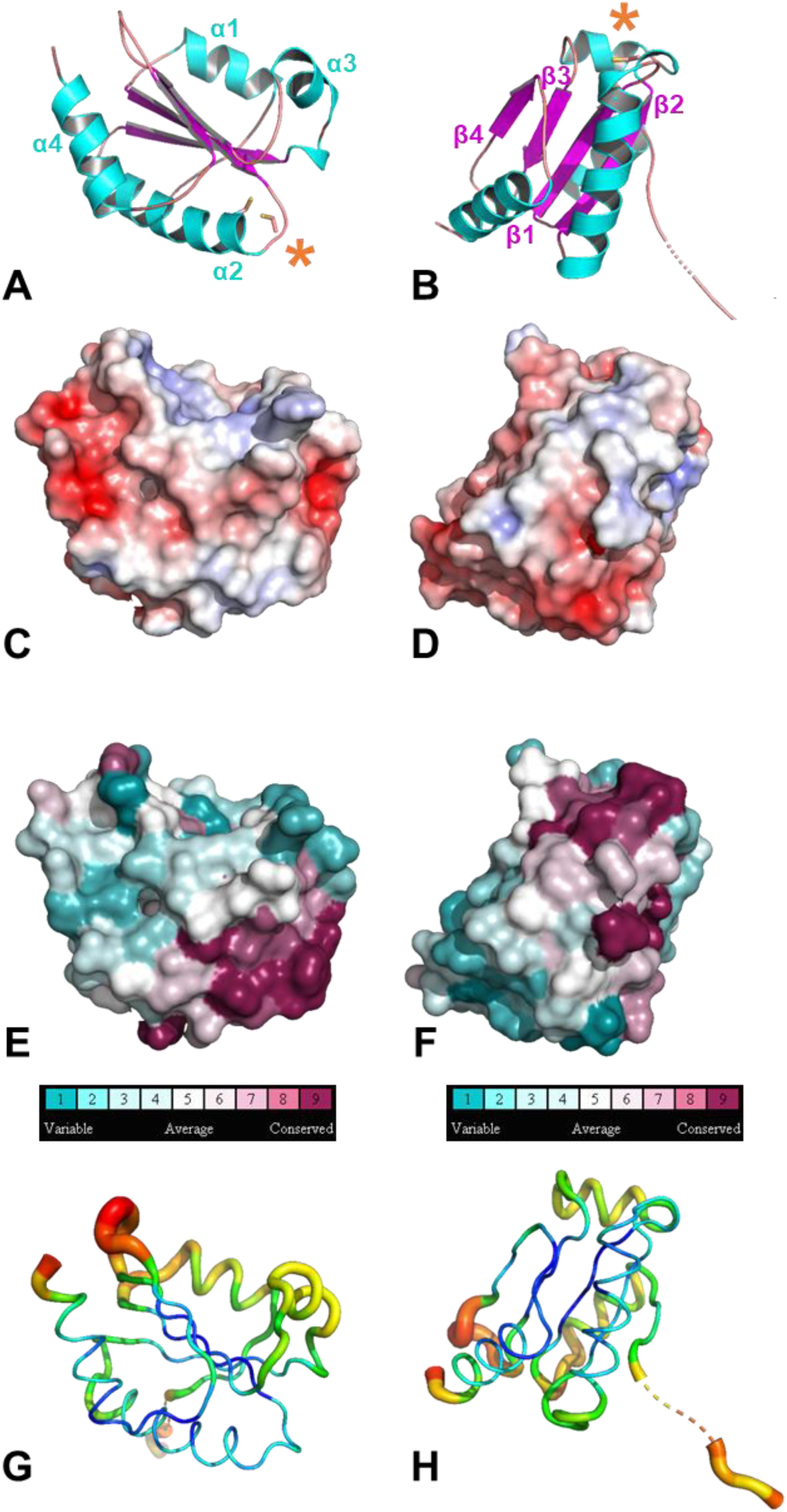
Crystallographic structure of chloroplastic CrTRXz. (A) Cartoon representation with helices 1-4 colored in cyan and strands 1-4 colored in magenta. Cysteines side chains are represented in sticks and highlighted with an asterisk. Residue numbering is in accordance with UniProt entry A8J0Q8-1. (B) 90° rotation around y-axis of A. (C) Electrostatic surface potential calculated with APBS [106]. Electronegative surface is colored in red while electropositive surface is colored in blue. (D) 90° rotation around y-axis of C. (E) Conservation of surface residues as calculated with CONSURF colored from purple (most conserved) to blue (least conserved). (F) 90° rotation around y-axis of E. (G) B-factor representation. High b-factor are colored in red and low b-factor are colored in blue.

### CrTRXz redox site and environment

Disulfide oxidoreductase activity of TRX relies on a pair of cysteines located at the amino-terminal tip of helix 2 in a conserved motif composed of the WCGPC peptide. The N-terminal active site cysteine (Cys_N_) attacks the disulfide bond on a target protein while the second active site cysteine termed Cys_C_ resolves the intermolecular disulfide bridge to free the reduced target protein and the oxidized TRX. CrTRXz mature sequence possesses two cysteines: Cys_N_ (C90) and Cys_C_ (C93) (Figure S1). Both are located at the tip of helix 2 (Figure 2A). Side chain thiols are modelled at 3.1 Å distance in what appears as a reduced state. Nucleophilic C90 displays a 13.56 Å^2^ solvent-accessible area, making it available for redox exchange with a target protein disulfide. The orientation of its thiol group is however unfavorable to thiol electron exchange because it points inwards in the direction of the TRX core. A local rearrangement of loop 88-91 would be required to tumble C90 thiol towards the solvent. Such a local rearrangement of TRX redox site would partly account for the entropy dependence of target disulfide binding [77]. In *Escherichia coli* TRX, D26 is buried in the core of the protein where it acts as a base catalyst in the thiol-disulfide interchange reaction [78, 79]. D31 was confirmed to play the same role in CrTRXh1, and this requires a water molecule for proton exchange [80]. CrTRXz also buries D84 at the equivalent position and its carboxyl side chain points in the direction of resolving C93 at a distance of 6.4 Å (Figure 2B). An intermediate water molecule, though not visible at the resolution of our structure, would fit in between at distances favoring hydrogen bonds with D84 and C93 (Figure 2B). P134 at the amino-terminal side of strand 3 faces the redox site at 3.5 Å and 4.6 Å of C93 and C90, respectively. P134 is built in the *cis* conformation. As reported earlier, this *cis*-proline is an important kinetic bottleneck for protein folding [81] and is already observed in inferred precambrian TRXs [82]. Altogether, redox site microenvironment and structural-dependent functional features of TRXs are present and canonical in CrTRXz.

**Figure 2.**
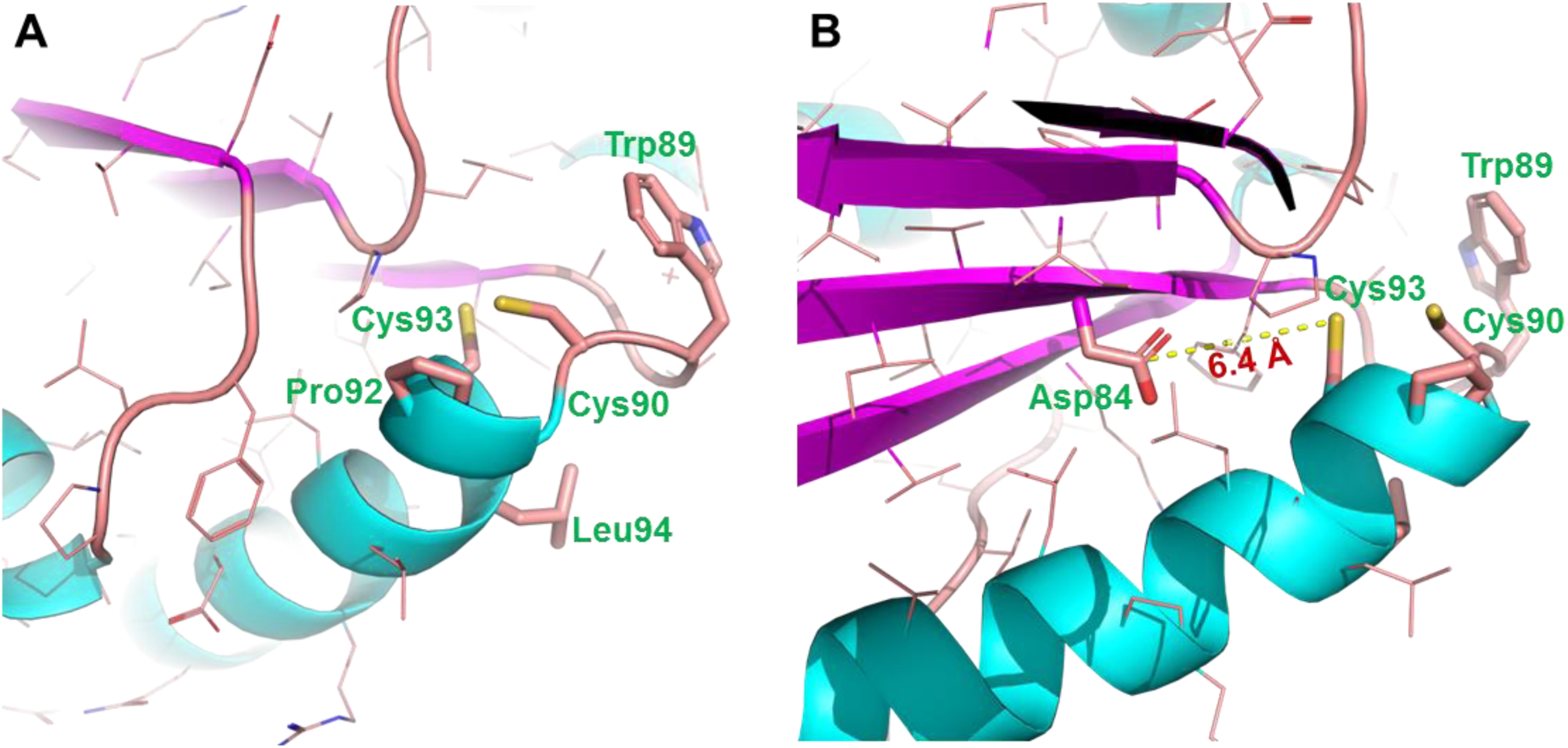
Redox site of CrTRXz. Main chain is represented as cartoon. Side chains are represented as lines and colored according to atom type (blue=nitrogen, red=oxygen, yellow=sulfur). (A) Redox site ^89^WCPGCL^94^ side chains are highlighted in sticks and labelled as three-letter code. C90 is located at the conserved nucleophile position. (B) Nucleophile D84 side chain is shown as sticks. The 6.4 Å carboxylate-thiol distance to C93 is drawn in broken line.

### CrTRXz quaternary structure

We unexpectedly observed early elution of purified CrTRXz from size-exclusion chromatography, showing an apparent higher molecular weight of the protein when compared to CrTRXf2. The experiment was repeated over two analytical gel filtration matrices (BioSEC3-300 and S6Increase) and the results were confirmed by comparison of distribution coefficients (Kav): Kav(CrTRXz)_BioSEC3-300_ = 0.625 ± 0.005, Kav(CrTRXf2)_BioSEC3-300_ = 0.89; Kav(CrTRXz)_S6Increase_ = 0.611, Kav(CrTRXf2)_S6Increase_ = 0.723 (Figure S3). According to BioSEC3-300 gel filtration calibrated with globular standards, the apparent molecular weight of CrTRXz is 30 ± 5 kDa, approximating twice the molecular weight of a 16 kDa subunit. Consequently, CrTRXz appears as a possible globular homodimer according to analytical size exclusion chromatography.

Analysis of protein interfaces in the crystal packing with PISA (EMBL-EBI) indeed uncovers six direct contact surfaces of 22.4 Å^2^, 98.9 Å^2^, 286.5 Å^2^, 322.1 Å^2^, 437.3 Å^2^ and 812.1 Å^2^ of the asymmetric unit monomer and its closest neighbors. The available surface of a monomer is 6848 Å^2^, hence the largest 812.1 Å^2^ interface covers 12% of CrTRXz surface. This crystallographic A:B dimer is formed by subunit A at positions x,y,z and subunit B at positions -x+2,-x+y+1,-z+2/3 in a neighboring unit cell (Figure 3A). PISA computes a solvation free energy gain (Δ^i^G) of −11.3 kcal.mol^-1^ upon formation of the interface. 21 residues are located at the interface. Ten residues form hydrogen bonds between R77_A_:E71_B_, R149_A_:M127_B_, E71_A_:R77_B_, M127_A_:R149_B_, and at equivalent yet non-symmetrical positions R129_A_:R129_B_. Four residues form multiple salt bridges between R77_A_:E71_B_ and equivalent E71_A_:R77_B_. Overall, PISA scores this interface with a complexation significance to 1 (stable assembly), while the other five detected interfaces score at 0 (no stable assembly).

**Figure 3.**
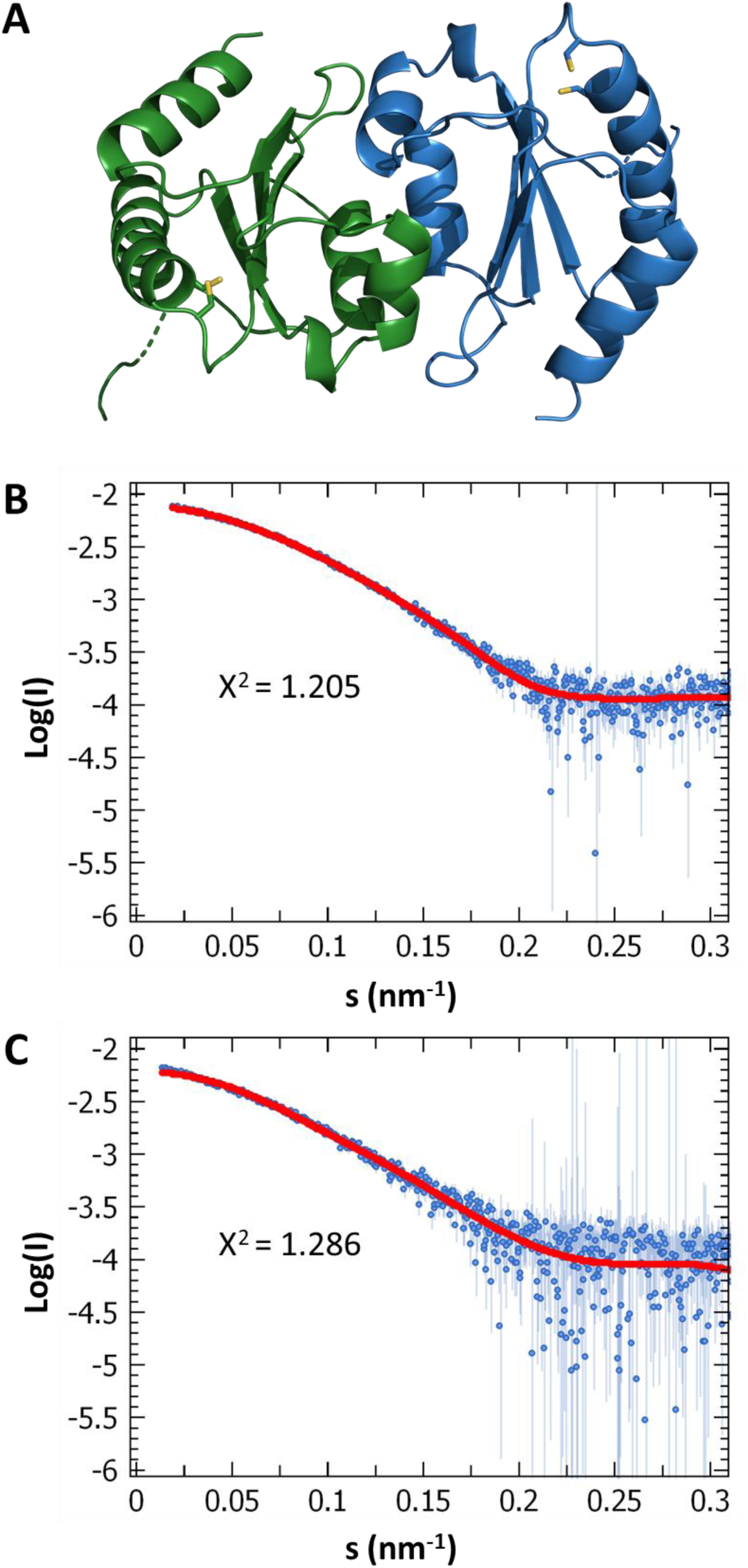
Thioredoxin z quaternary structure. (A) Crystallographic dimer of CrTRXz represented in cartoon with reactive cysteines side chain in sticks. Chain A is colored in blue, chain B in green. (B) SAXS scattering curve log(I)=f(s) of batch 1 of CrTRXz in blue was fitted to the crystallographic monomeric model of CrTRXz in red. (C) Same experiment as B. with batch 2 of CrTRXz.

To try and probe the quaternary structure of CrTRXz in solution, we submitted two independent preparations of pure CrTRXz to small-angle X-rays scattering coupled to size-exclusion chromatography (SEC-SAXS) (Figure 3B,C). Guinier analyses radii of gyration of 20.16 ± 0.25 Å (protein preparation 1) and 23.34 ± 0.43 Å (preparation 2) allowed computation of respective molecular weights intervals of 15.80-17.75 kDa and 18.35-20.85 kDa at a credibility probability higher than 95%. The oligomeric state of CrTRXz detected by SAXS is consequently that of a monomer of 16.2 kDa. Analysis of CrTRXz by mass spectrometry under native or denaturant conditions showed only the presence of monomeric species in our experimental conditions suggesting that in solution CrTRXz is mainly monomeric (data not shown).

In conclusion, the dimerization observed in crystal packing is too weak or too transitory to represent the principal protein population *in vitro* in the SAXS and MS experimental setups, while the higher apparent molecular weight on size exclusion chromatography may be due to non-globularity. The monomeric state is the most probable form expected in the diluted, non-crystalline physiological context.

### CrTRXz reactive surface displays a unique electronegative potential

Multiplicity of TRX types in the chloroplast stroma raises the question of their specificity and the underlying physico-chemical determinants for selective and non-overlapping disulfide bond recognition in different targets [40]. In order to identify z-type specificities, we analyzed the high-resolution crystal structure of CrTRXz. The secondary structure topology is almost identical to other known TRXs (Figure S2). The redox residues are also positioned at the canonical sites and they form the most conserved surface of the protein among homologs (Figure 1E and 1F) [83]. The dynamics of the main chain is approximated by the crystallographic b-factors, the highest values of which are grouped at the amino-and carboxy-terminal ends and at the loop connecting strands 3 and 4 (Figure 1G and 1H). Helices 3 and 1 also group higher than average b-factors, albeit to a lower extent than the aforementioned loop and N/C-termini. This local flexibility may predict functional deformations of CrTRXz structure upon folding, interaction with protein partners, or during allosteric controls. All dynamic elements are however located far from the redox site and they are unlikely to contribute alone to the selectivity of CrTRXz towards targets.

The principal determinant of CrTRXf2 selectivity was described as an electropositive crown of residues exposed at the surface surrounding the redox site [55–57, 84]. We compared the electrostatic surface of CrTRXz and observed large electronegative patches, and a general lack of electropositive elements (Figure 1C and 1D). Notably, CrTRXz presents the z-type conserved apolar L94 immediately after the redox site WCGPC, while all other plastidial TRX possess in this position either a positively charged residue (R/K for m-, x- and f-type) or an hydrogen bond donor residue (Q for y-type). Differential electrostatic repartition is particularly marked in the vicinity of the redox site and represents a significant qualitative difference with CrTRXf2 and CrTRXm equivalent surfaces (Figure 4B, 4D and 4F). The principal determinants of CrTRXf2 selectivity were proposed to be K78, K114, N130, K131, N133, K134, K143 and K164 (PDB entry 6I1C numbering [57]). Equivalent positions on structurally aligned CrTRXz are occupied by E63, Q102, D119, E120, N121, P122, Q131 and E151. Seven out of these eight residues are conserved in TRXz homologs (Figure S1), supporting an important function for type-z physiological functions in plants. In both CrTRXf2 and CrTRXz crystal structures, the aforementioned side chains are located at the protein surfaces and point their polar groups to the solvent. If the amine group of lysines expose a +1 charge while the carboxyl group of aspartates and glutamates expose a −1 charge at physiological pH, aforementioned CrTRXf2 selectivity surface will accumulate a net charge of +6 while corresponding positions on CrTRXz sum a net charge of −4. The contributions of asparagine and glutamine residues to each selectivity surfaces are less explicit since they potentially behave as hydrogen bond donors or acceptors.

**Figure 4.**
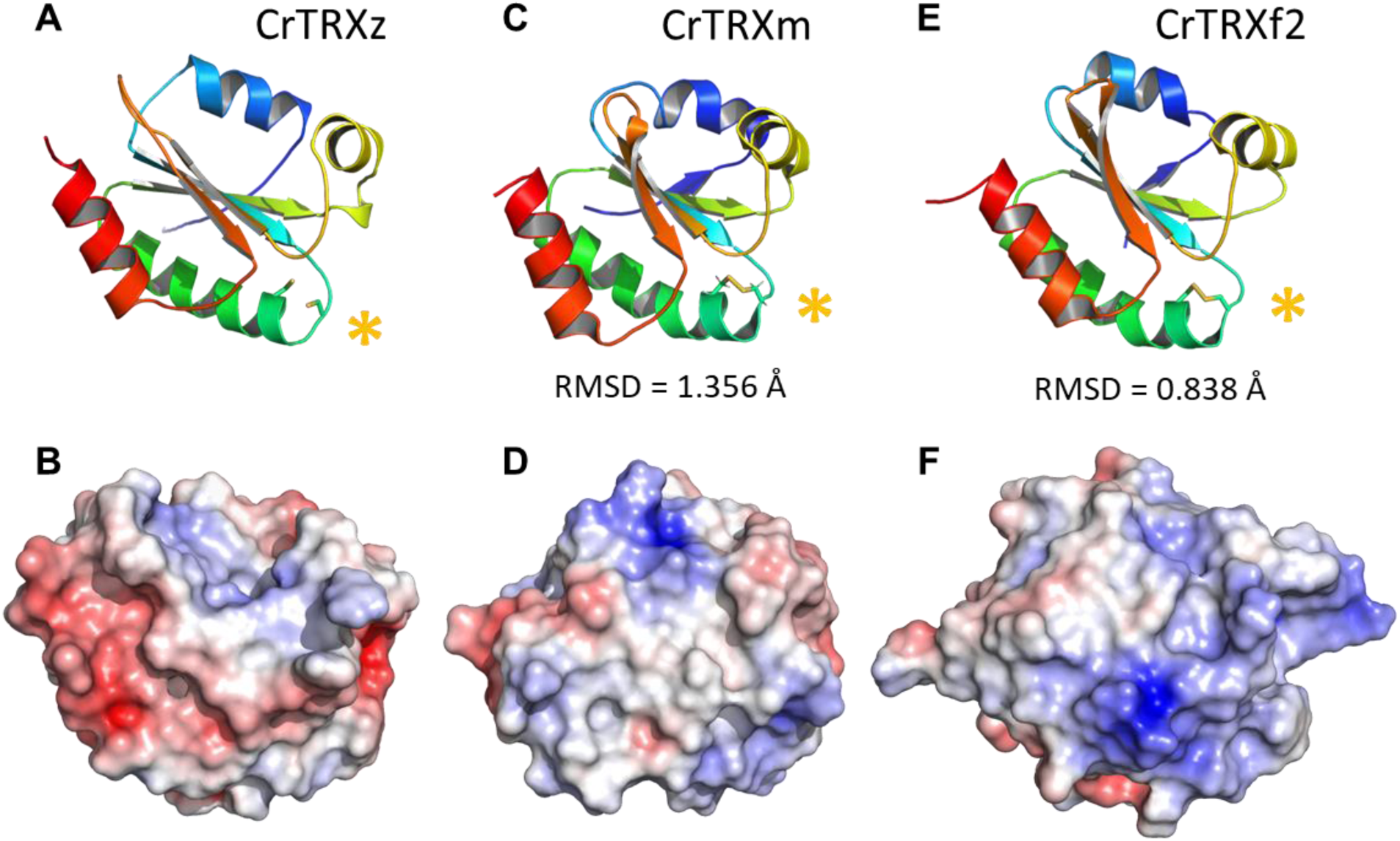
Structural comparison of plastidial TRX experimental structures. Root mean square deviations (RMSD) between equivalent alpha carbons were calculated with PYMOL. (A) CrTRXz crystal structure (this study) represented in cartoon and colored from blue (amino-terminus) to red (amino-terminus). Cysteines sidechain are shown in sticks representation and highlighted with an asterisk. (B) Electrostatic surface potential of CrTRXz calculated with APBS [106]. Electronegative surface is colored in red while electropositive surface is colored in blue. (C) CrTRXm nuclear magnetic resonance structure[107] (PDB identifier 1DBY) represented and colored as in A. (D) Electrostatic surface potential of CrTRXm calculated with APBS and colored as in B. (E) CrTRXf2 crystal structure (PDB entry 6I1C) represented and colored as in A. (F) Electrostatic surface potential calculated with APBS of CrTRXf2 and colored as in B.

Overall, TRXz and TRXf2 recognition modes probably rely on distinct surface properties of targets because they present opposite electrostatic profiles. According to their mature sequence, CrTRXz presents an acidic isoelectric point while that of CrTRXf2 is basic (4.55 and 8.45, respectively). CrTRXm also presents an acidic isoelectric point (5.09), but important surface differences with CrTRXz stem from the charges repartition around the redox surface (Figures 4B and 4D).

### TRXz is electro-complementary to known protein interactants

Four candidate proteins are described as TRXz interactants in *Arabidopsis thaliana*: FLN1, FLN2, PRIN2 and MRL7. The molecular structure of these proteins is not experimentally determined but their structures were predicted by homology using the PHYRE software [85] (Figure 5). AtFLN1 possesses five cysteines and based on the protein model, adjacent C105 and C106 are buried within the structure and seems to be affected by TRX only in a solvent exposed unfolded state. C204, C398 and C418 are predicted to be solvent-exposed and available for TRX redox interaction (Figure 5A) even though they probably do not form intramolecular disulfide bridges as they appear too distant. C398 is surrounded by an electropositive surface complementary to CrTRXz recognition surface (Figure 5B). Homology-modelled AtFLN2 includes nine cysteines, consecutive C208 and C209 are also located under the surface and probably not accessible in the folded state (Figure 5C). C290, C307, C322, C326, C419 are present at the predicted surface while C529 hidden to solvent and C253 is at the bottom of a deep and narrow cleft. C290, C322, C326 and C327 are surrounded by electropositive surfaces that qualifies them as potential targets of TRXz redox interaction (Figure 5D). The three later would make good candidates for intra-molecular dithiol/disulfide exchange since their thiols are predicted at 7.7, 8.5 and 9.2 Å distances. AtMRL7 model encompasses two cysteines, both predicted at the solvent exposed surface of the protein (Figure 5E). C281 is surrounded by marked electropositive potentials and may then be targeted by electro-complementary TRXz redox surface (Figure 5F). AtPRIN2 could not be modelled by PHYRE as a globular structure, because of the lack of close homology with experimental models (best sequence identity 27 % over 32 % residue coverage, or 22 % identity for 49 % coverage). However, the 20-residues peptide NH_3_^+^-LSSLSRRGFV<u>C</u>RAAEYKFPDP-COO^-^ centered on redox target C68 (underlined) contains three arginines and one lysine and even though the mature protein sequence confers an acidic pI (4.43), the local environment of the cysteine in this peptide presents a basic pI of 9.31 that would make it a reasonable target for interaction with TRXz electronegative surface.

**Figure 5.**
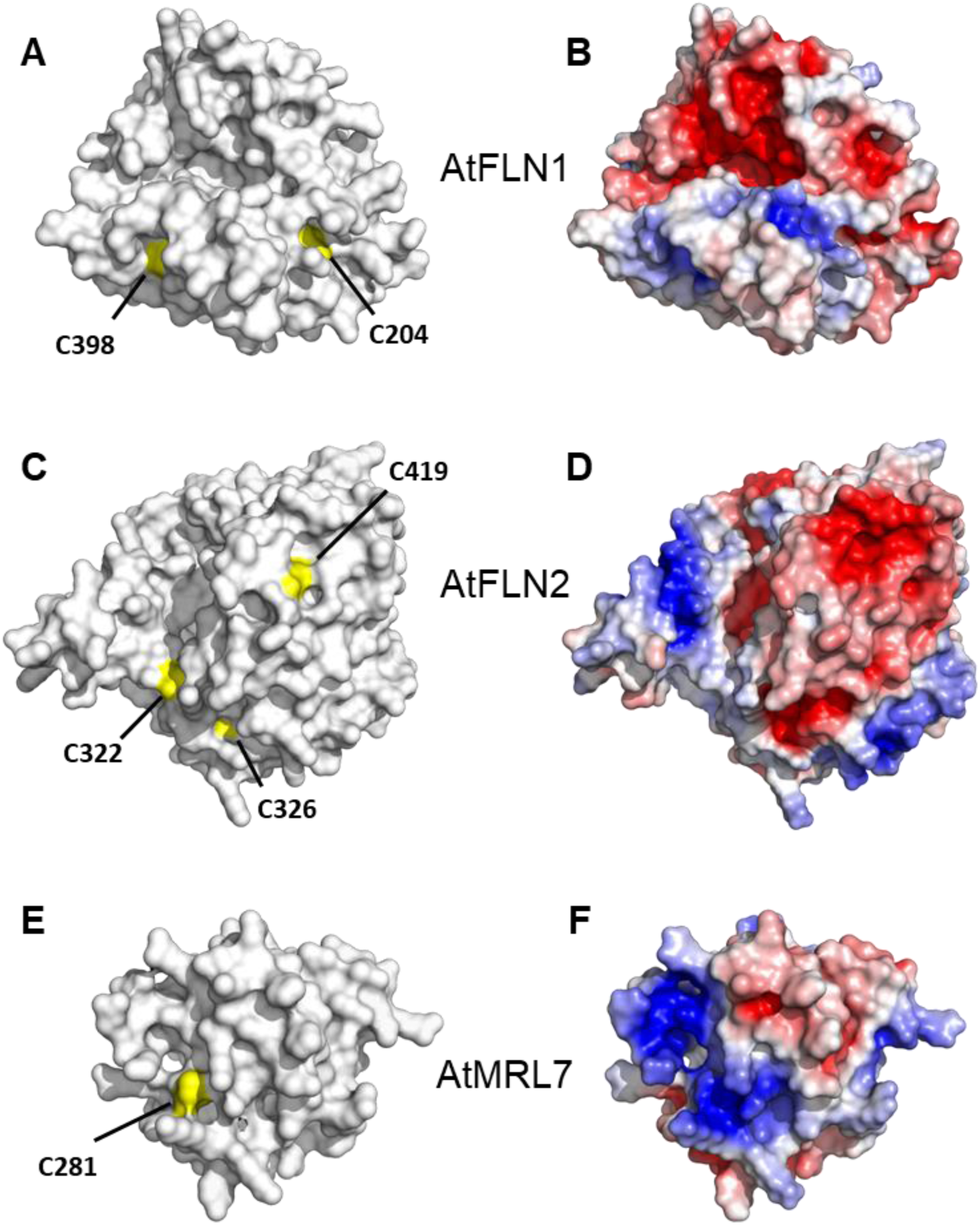
Electrostatic surface of PHYRE2 [85] homology-modelled TRXz protein partners. (A) Connolly solvent exclusion surface of AtFLN1 residues 90-464 from the sequence of UniProt entry Q9M394 with Cysteines in yellow. (B) Electrostatic surface potential calculated with APBS. Electronegative surface is colored in red while electropositive surface is colored in blue (C) Connolly solvent exclusion surface of AtFLN2 205-557 from the sequence of UniProt entry F4I0K2 with Cysteines in yellow. (D) Electrostatic surface potential calculated with APBS and colored as in B. (E) Connolly solvent exclusion surface of AtMTR7 residues 195-312 from the sequence of UniProt entry F4JLC1 with Cysteines in yellow. (F) Electrostatic surface potential calculated with APBS and colored as in A.

### *CrTRXz can activate CrPRK* in vitro

While CrPRK is a well-known CrTRXf2 target, presumably because of its partially negative surface around C55, another large positive patch is also present near C55 at the enzymatic pocket [48, 54]. The mixed environment of CrPRK C55 suggests an independent, alternative mechanism of redox activation by TRXz. We probed the activation efficiency of the CrTRXf2, CrTRXz, and CrTRXm on PRK activity. The ability of CrTRXs to reductively activate oxidized CrPRK was measured using different TRX:PRK ratios and assaying the enzyme activity after 1 h of incubation (Figure 6). In every incubation, CrPRK was 5 μM and TRX varies from 0.5 μM (ratio 1:10 TRX:PRK) to 50 μM (ratio 10:1 TRX:PRK). At the lowest ratio (1:10) no TRX significantly activated PRK, while at higher ratios both CrTRXz and CrTRXf2 activated the enzyme. The CrTRXz activation plateaued at 2-fold excess, reaching 60% of maximal activity. No further activation was observed at higher TRX:PRX molar ratios. PRK activity in the presence of CrTRXf2 increased until 6-fold excess, where 100% of maximal activity was obtained. CrTRXm was able to partially activate PRK only at the highest molar ratios (*i*.*e*. 6:1 and 10:1 TRX:PRK ratios). These results demonstrate that CrTRXf2 is the main activator for PRK *in vitro*, whereas CrTRXz is able to activate CrPRK to a lesser extent.

**Figure 6.**
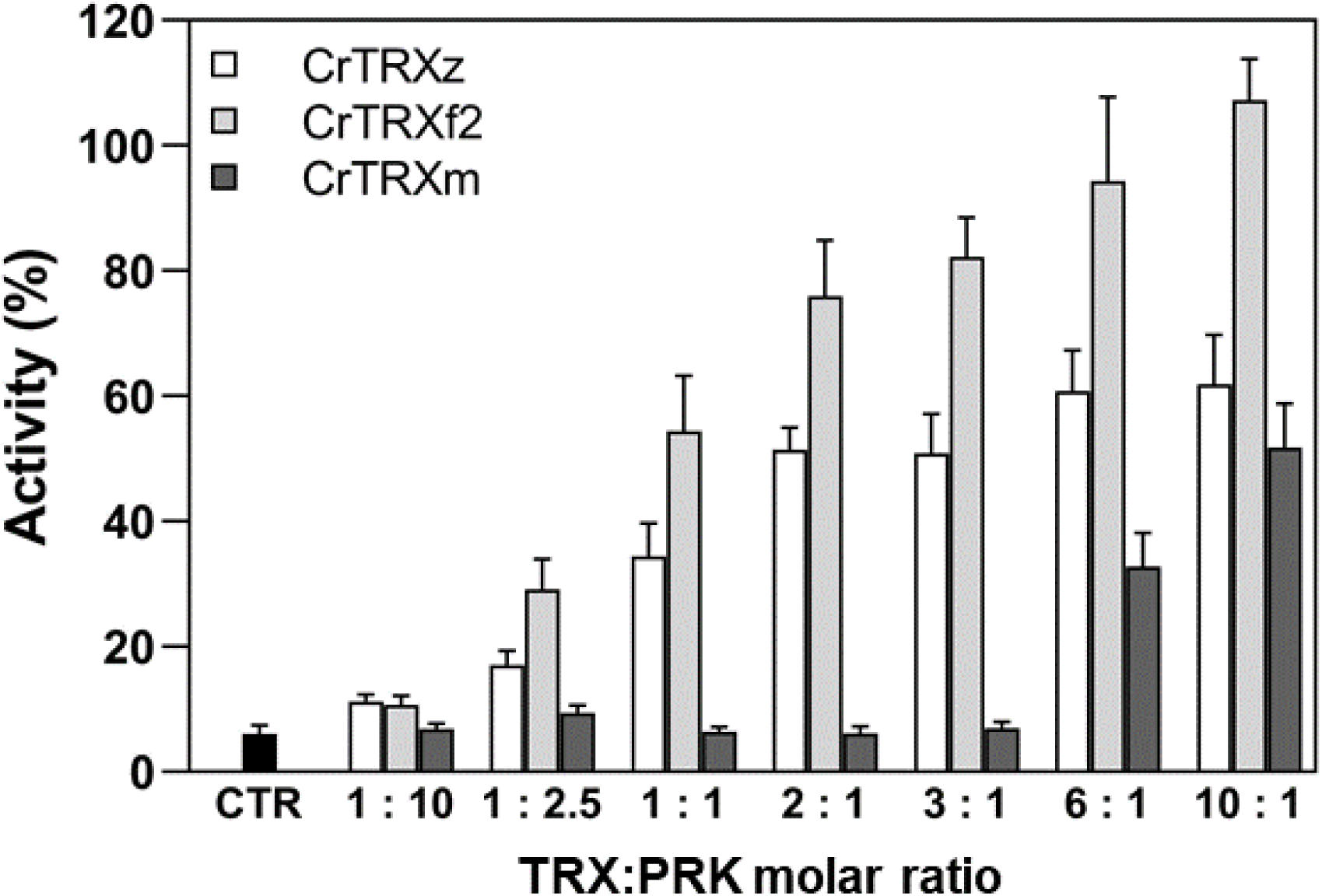
Activation efficiency of CrTRXz, CrTRXf2, and CrTRXm on CrPRK. Activation of oxidized CrPRK (5 μM) after 1 h incubation with increasing concentrations (0.5-50 μM) of CrTRXz (white bars), CrTRXf2 (light grey bars) and CrTRXm (dark grey bars) in presence of 0.2 mM DTT. TRX:PRK molar ratios (n:n) are indicated for each tested condition. Control sample (CTR, black bar) shows the activity after incubation with 0.2 mM DTT without TRXs. All the activities were normalized on fully activated PRK obtained after 4 h of reduction with 40 mM DTT. Data are reported as mean ± standard deviation (n=3).

## Discussion

Here, we report the first experimental structure of a TRXz from the model eukaryotic alga *Chlamydomonas reinhardtii* (PDB ID code: 7ASW). Structural analysis reveals that chloroplastic CrTRXz is a classical TRX with an atypical electronegative surface conserved in plant homologs. We dissected TRXz surface properties to assess its putative specificity over known target proteins of AtTRXz. Moreover, based on structural features, we proposed the Calvin-Benson enzyme CrPRK as being regulated by CrTRXz, which was confirmed by our *in vitro* activity assays.

### CrTRXz is a canonical thioredoxin

The determination of the crystal structure of chloroplastic TRXz from the green alga *Chlamydomonas reinhardtii* confirmed that it folds as a canonical αβα compact domain exposing the conserved WCGPC redox pentapeptide at nucleophile attacking distance of incoming target disulfide bond. Other signatures of reactivity and folding were observed in our structure, *e*.*g*. the base catalyst aspartate in a water-bridging distance to resolving Cys_C_, the ten-residue helix 1 of eukaryotic TRXs and the folding bottleneck *cis*-P134. The reactive Cys redox pair appears dissociated in a dithiol state available for the reduction of a target disulfide bond.

### CrTRXz evolved distinct features

Although conform to the thioredoxin standard, our crystal structure reveals distinct features on CrTRXz. The peptide connecting strand 2 and helix 3 is folded as a short helix 3_10_ that is stabilized by hydrogen-bonding interaction of D119 with W89, adjacent to Cys_N_ in the redox site. This particular state may modify the reactivity of TRXz active site or rather a functional conformation of TRX since D119 is conserved in CrTRXx, y, h1, h2, and o while it is substituted by an isosteric asparagine in CrTRXf2 and m. C126 of AtTRXf1 was shown to be glutathionylated and the glutathionylation demonstrated to subsequently affect the redox capacity of this TRX [86]. CrTRXf1 and f2 but not the other TRX types possess this regulatory cysteine, at the first position after aforementioned D119 and its equivalents. Hence, regulation of thioredoxin by redox signaling may be caused by distal effects on the redox site through a bonding network similar to CrTRXz D119-W89. Interestingly, selective inactivation of TRXf by glutathionylation would leave TRXz as an alternative to perform photosynthetic functions such as light- and TRX-dependent activations of PRK.

CrTRXz crystallizes as a homodimer and elutes as larger than globular monomer objects as judged by size-exclusion chromatography. Small-angle X-rays scattering (SAXS) and native mass spectrometry however unequivocally contradicted the dimer hypsothesis in solution, with the measured gyration radii and diffusion curves revealing a monomeric state of globular CrTRXz *in vitro*. Dimerization is either only a crystallization artifact or a transitory interaction that controls CrTRXz function in restricted conditions of high local concentration, like those of a liquid-liquid phase separation or of a multiprotein complex. The interaction surface is distant from the redox site and may rather serve to scaffold TRXz-containing complexes with internal two-fold symmetries. Other TRX quaternary states were tested in the past and proved monomeric [87, 88] in reduced state [89].

### Electronegativity as a driver of CrTRXz specific complexation

Western blot protein quantification of *Arabidopsis thaliana* leaf extracts indicated relative abundances of 69.1% TRXm, 22.2% TRXf, 6.3% TRXx, 1.3% TRXy and 1.1% TRXz, representing a TRXz concentration of 1-5 pmol.mg^-1^ stromal protein [90]. Notwithstanding its relative scarcity, TRXz still makes significant contributions to chloroplast functions as demonstrated by the lethality of its deletion mutant. We propose that TRXz loss cannot be compensated by other TRXs because of the irreplaceable specialization of its electronegative surface. Described determinants of TRXf2 selectivity present an opposite, electropositive character compared to TRXz. Reported targets of TRXz in *Arabidopsis thaliana* present exposed cysteines on electropositive surfaces that match TRXz redox site (Figure 5). One interesting candidate for interaction is TRXf itself, since the electropositive environment of its active site is an ideal electro-complementary platform for TRXz docking. Consistently, TRXz was reported to be reduced *in vitro* by FTR [67] but also by other TRX types [69] suggesting that it could play a specific role in reductive or oxidant signaling [64, 69].

We report here that CrTRXz is able to activate CrPRK, a known target of the main CBC activator TRXf2. CrTRXz activation is slightly weaker than that of CrTRXf2 but stronger than CrTRXm activation. It relies on the ratio between TRX and target, so TRXz could efficiently activate PRK or other targets when it reaches a certain concentration in the chloroplast. Interestingly, the cell has at its disposal two different TRXs to activate PRK and tune the CBC flux according to environmental conditions. In conditions when TRXf2 is inactivated by glutathionylation [86], one can hypothesize that TRXz becomes the replacement activator of PRK.

TRX complementarities should now be confirmed by the description of the actual complexes between TRX and targets, ideally from experimental structures at atomic resolution. TRXm, x, and y in the chloroplast still await a comprehensive description of the molecular basis for specific disulfide targeting, as well as TRXh/TRXo and bacterial TRX in their respective compartments. Later on, a global simulation of electron transfers in the proteome will require knowledge on the concentrations of all TRXs, reductant sources, and targets, their mutual binding affinities, interaction turnovers and their cysteine redox potentials.

## Material and methods

### Cloning

Nuclear encoded amino acid sequence of *Chlamydomonas reinhardtii* thioredoxin z (UniRef100 entry A8J0Q8, thioredoxin-related protein CITRX) was analyzed by TargetP2.0 [91, 92], ChloroP [93] and Predalgo [94] to predict the transit peptide cutting site and the subsequent mature sequence of chloroplastic protein. The sequence coding for amino acids 56-183 was PCR-amplified from *Chlamydomonas reinhardtii* EST database plasmid AV642845 with DNA primers 5’-TGTCGC**CATATG**GTCATTAGCCATGGAAAG-3’ and 5’-CTCCTT**AAGCTT**CTACTGCTGCGGCGCCTCGGG-3’. The PCR product was inserted into pET28a by restriction with NdeI and HindIII and subsequent ligation, yielding pET28a-His_6_-CrTRXz. Recombinant protein is a fusion of Met-HHHHHH-SSFLVPRGSHM- and TRXz residues 56 to 183, as validated by plasmid Sanger sequencing. Residue numbering is accorded to UniProt sequence A8J0Q8-1 throughout the text.

### Expression and purification

*Escherichia coli* strain BL21(DE3) Rosetta-2 pLysS (Novagen Merck, Darmstadt Germany) was transformed with pET28a-His6-CrTRXz and grown in 1 L of 2YT medium supplemented with kanamycin (50 µg.mL^-1^) and chloramphenicol (34 µg.mL^-1^) to exponential phase at an optical density of 0.6 before T7-RNA polymerase induction of overexpression with 200 µmol.L^-1^ IPTG for 3.5 h at 37°C. 4.1 g of cell pellet were resuspended in 20 mL buffer A (20 mmol.L^-1^ Tris-Cl pH 7.9, 200 mmol.L^-1^ NaCl), lysed by 2 min sonication with 0.4 sec pulses at output 5 on W-375 sonicator equipped with microtip (Qsonica, Newtown CT USA) on ice. Total extract was clarified by 20 min centrifugation at 30,000 rcf at 4°C and the soluble fraction was loaded on affinity chromatography over 5 mL of NiNTA resin (Sigma-Aldrich Merck, Darmstad Germany). Resin was step-washed with 10 mL of buffer A supplemented with 10, 20, 30, 40 or 50 mmol.L^-1^ imidazole. Elution was executed with steps of 10 mL of buffer A supplemented with 100, 150, 200 or 250 mmol.L^-1^ imidazole. Collected fractions were analyzed by Coomassie stained SDS-PAGE that revealed the presence of >99 % pure TRXz in wash fractions 20, 30, 40 and 50 and elution fractions 100 and 200. Pooled fractions were buffer exchanged to buffer A and concentrated by ultrafiltration on 3000 MWCO Amicon filter units (Millipore Merck, Darmstadt Germany). A final concentration of 10.2 mg.mL^-1^ was measured by NanoDrop 2000 spectrophotometer (Thermo Fisher Scientific, Waltham MA USA) with theoretical Mw = 16,191.5 g.mol^-1^ and ε_280_ = 5,567.5 mol^-1^.L.cm^-1^.

### Size-exclusion chromatography

0.1 mg affinity-purified recombinant CrTRXz was loaded on Superose6 10/300 Increase column or BioSEC3-300 column in 20 mmol L^-1^ Tris-Cl pH=7.9, 100 mmol.L^-1^ NaCl. Isocratic elution was recorded at 280 nm.

### Small angle X-rays scattering

Diffusion curves were collected from BioSEC3-300 size-exclusion chromatography in line with X-ray exposure capillary at beamline SWING of synchrotron SOLEIL (Saint-Aubin, France). Data were processed with FOXTROT (Xenocs, Grenoble France) using CLEVERAVG algorithms and interpreted with ATSAS [95, 96].

### Crystallization and structure determination

Purified CrTRXz was tested for crystallization on commercial sparse-screening conditions (Qiagen, Hilden Germany) of the Joint Center for Structural Genomics screens [97] with a mixture of 30 nL protein and 30 nL precipitant equilibrated against a reservoir volume of 30 µL mother liquor. 30 µm sided monocrystals grew in one week in condition 38 at position D2 of screen IV (0.1 mol.L^-1^ HEPES pH=7,5; 1,26 mol.L^-1^ ammonium sulfate). Crystals were transferred in mother liquor supplemented with 12.5% glycerol and flash-frozen in liquid nitrogen for diffraction experiments at micro-focused beamline PROXIMA-2A at SOLEIL synchrotron (Saint-Aubin, France). A 99.36 % complete dataset was collected at 2.444 Å resolution. Data was indexed in space group *P*3_2_21, integrated, scaled and converted with XDSME [98]. Structure was phased by molecular replacement with PHENIX [99, 100] PHASER-MR [101] using CrTRXf2 (PDB 6I1C [57]) as a search model and a single protein in the asymmetric unit. Model was refined by iterative cycles of manual building in COOT [102, 103] followed by refinement with PHENIX.REFINE [104] until completion of a structure passing MOLPROBITY [105] evaluation with 100 % residues in Ramachandran restrains, RMS(bonds)=0.010, RMS(angles)=1.44 and final R_work_=0.2296, R_free_=0.2335 (Table 1). Structure representations were drawn with PYMOL (Schrodinger, New York USA).

### Thioredoxin activation assay

*Chlamydomonas reinhardtii* phosphoribulokinase (CrPRK) was recombinantly expressed as published earlier [48]. Pure CrPRK was oxidized in presence of 0.2 mM 5,5’-dithiobis(2-nitrobenzoic acid) (DTNB, Sigma-Aldrich) for 45 min at room temperature. CrPRK was desalted to stop the oxidation and used to assay TRX activation efficiency. Oxidized CrPRK at 5 μM was incubated in presence of 0.2 mM dithiothreithol (DTT) and from 0.5 μM to 50 μM CrTRXz, CrTRXf2 and CrTRXm for 60 minutes, then PRK activity was assayed as in [48]. Control activity was obtained incubating CrPRK with 0.2 mM DTT for 60 min. Activities were normalized on fully reduced PRK obtained after 4 h incubation at room temperature in presence of 40 mM DTT.

### Structural data

Validated crystallographic reflection data and model are deposited in the protein data bank under access code 7ASW.

## Acknowledgements

This work was funded by CNRS, Sorbonne Université, and Agence Nationale de la Recherche grants LABEX DYNAMO 11-LABX-0011, CALVINDESIGN (ANR-17-CE05-001) and CALVINTERACT (ANR-19-CE11-0009). Libero Gurrieri was supported by grant PON ARS01 00881 “ORIGAMI” (Italian Ministry of University and Research). Fernanda Pires Borrega and Sandrine Hoang contributed to the preparation of recombinant thioredoxin z. The *Institut de Biologie Physico-Chimique* (FR 550 CNRS) provided access to crystallization facilities. We acknowledge SOLEIL for provision of synchrotron radiation facilities. We thank Martin Savko, Serena Sirigu, and William Shepard for assistance in using beamline PROXIMA-2a, and Aurélien Thureau and Javier Perez for assistance in using beamline SWING.

## Supplementary figures

**Supplementary figure 1.**
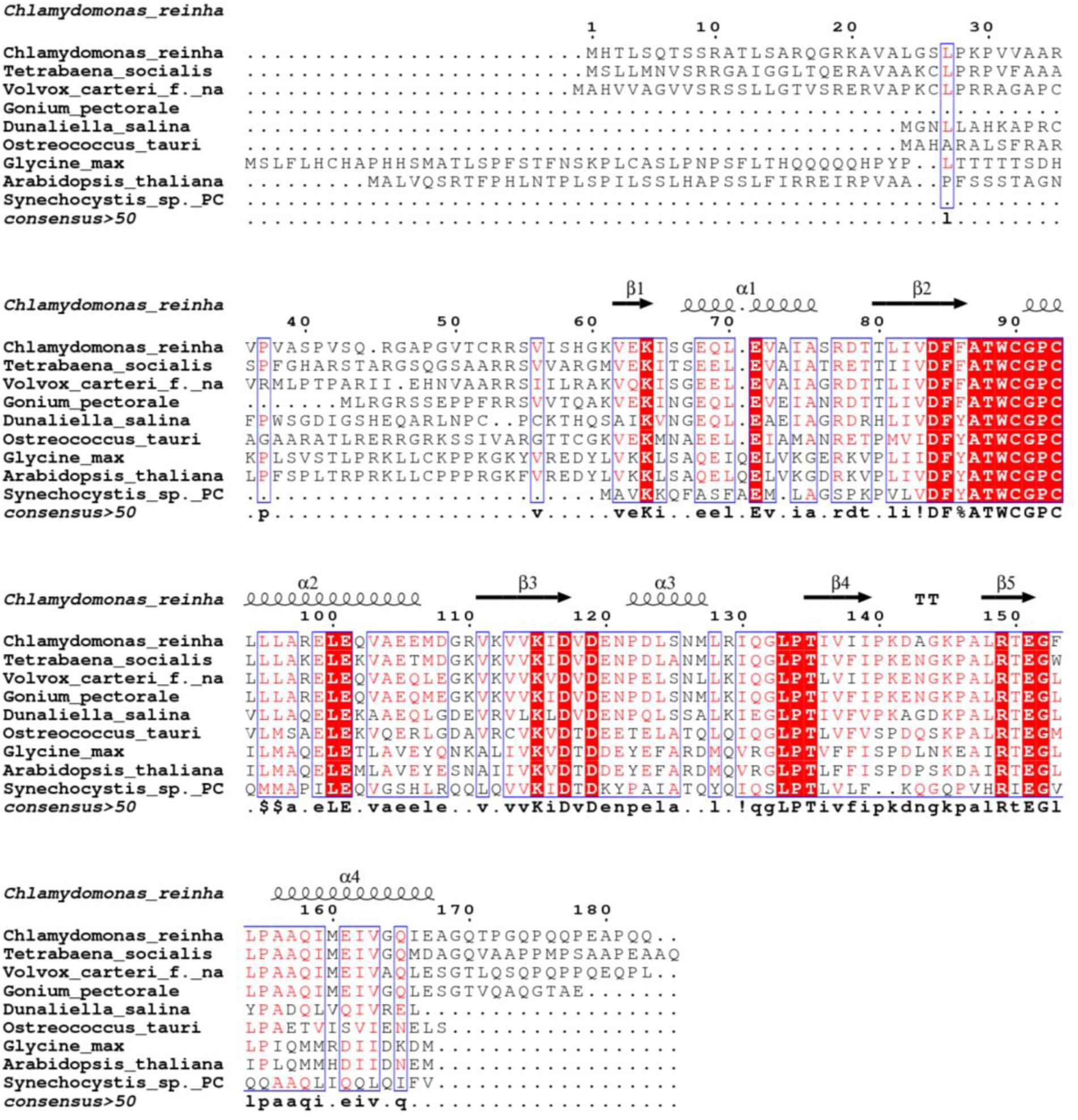
Sequence alignment of CrTRXz homologs. Sequence alignment was performed by Clustal Omega with homologs from different species. Residues with more than 50% conservation are colored in red, strictly conserved residues are in white with a red background. Numbering is according to the sequence of TRXz from *Chlamydomonas reinhardtii*. TRX sequences were retrieved from UniprotKB database for the following species: *Chlamydomonas reinhardtii* (A8J0Q8), *Gonium pectorale* (A0A150FZQ6), *Tetrabaena socialis* (A0A2J7ZQ73), *Volvox carteri f. nagariensis* (D8TVT1), *Ostreococcus tauri* (A0A1Y5IIS5), *Arabidopsis thaliana* (Q9M7×9), *Synechocystis sp. PCC 6714* (A0A068MSG5), *Glycine max* (I1JW39).

**Supplementary figure 2.**
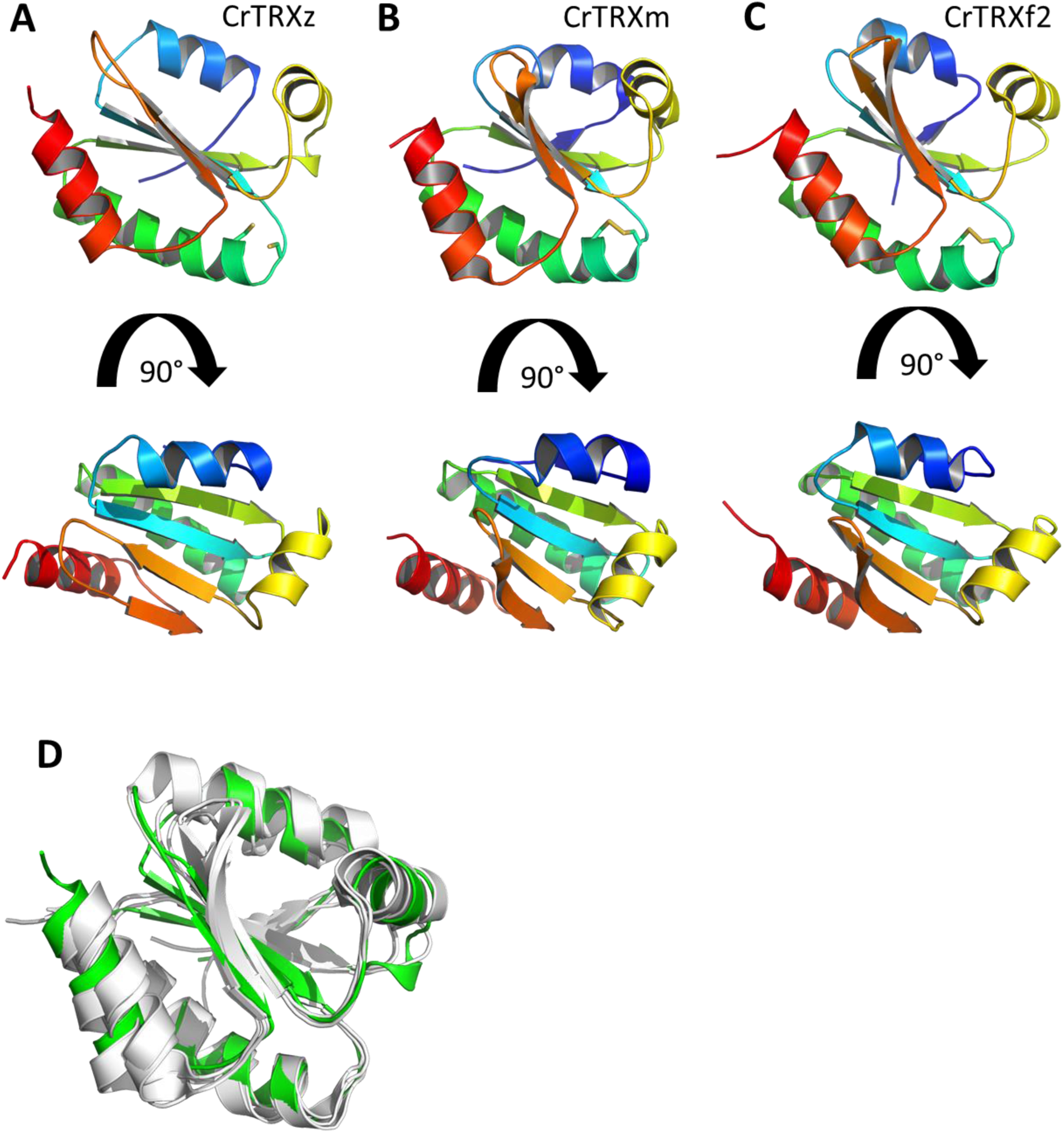
CrTRX experimental structures. (A) Side and top view of CrTRXz structure. CrTRXz is represented in cartoon colored in rainbow from N-terminus (blue) to C-terminus (red). Reactive cysteine side chains are represented in sticks. (B) CrTRXm in the same orientation, representation and coloration as A. (C) CrTRXf2 in the same orientation, representation and coloration as A. (D) Structural alignment of CrTRXf2, CrTRXm and CrTRXh structure colored in white on CrTRXz colored in green structure realized by PyMOL. RMSD of alignments are respectively 1.357 Å, 0.838 Å and 1.123 Å for CrTRXm, CrTRXf2 and CrTRXh.

**Supplementary figure 3.**
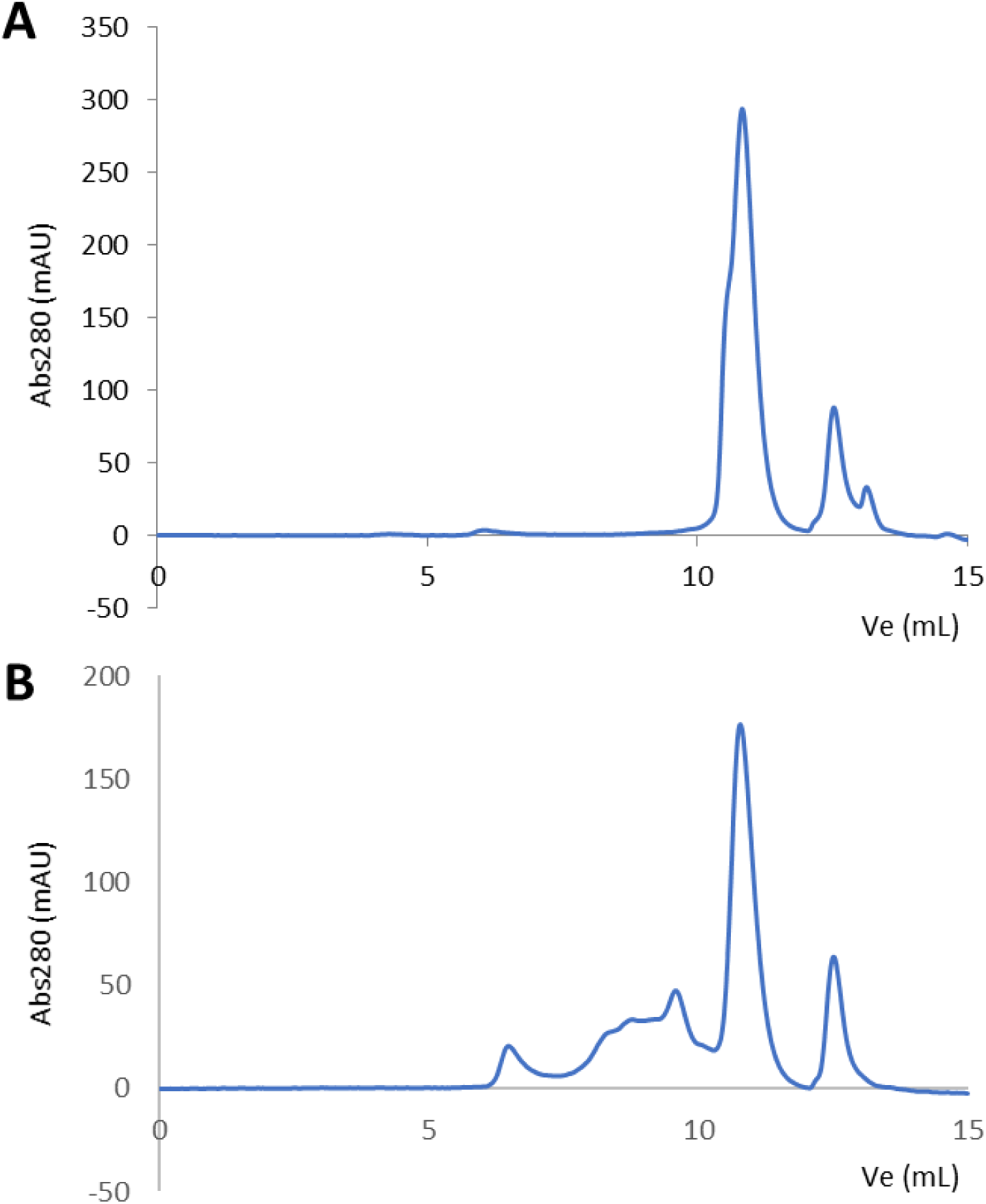
CrTRXz size-exclusion chromatography. High-pressure liquid chromatography profile over BioSEC3-300 column of (A) preparation 1 and (B) preparation 2 of CrTRZx. Globular calibration standards elute at 7.2 mL (670 kDa), 9.2 mL (158 kDa), 10.6 mL (44 kDa), 11.7 mL (17 kDa). Dead column volume is 6.5 mL and total column volume is 13.4 mL.

